# iDamage: a method to integrate modified DNA into the yeast genome

**DOI:** 10.1101/510966

**Authors:** Katarzyna H. Masłowska, Luisa Laureti, Vincent Pagès

## Abstract

In order to explore the mechanisms employed by living cells to deal with DNA alterations, we have developed a method by which we insert a modified DNA into a specific site of the yeast genome. This is achieved by the site-specific integration of a modified plasmid at a chosen locus of the genome of *Saccharomyces cerevisiae*, through the use of the Cre/*lox* recombination system. In the present work, we have used our method to insert a single UV lesion into the yeast genome, and studied how the balance between error-free and error-prone lesion bypass is regulated. We show that the inhibition of homologous recombination, either directly (by the inactivation of Rad51 recombinase) or through its control by prevenAng the poly-ubiquitination of PCNA (*ubc13* mutant), leads to a strong increase in the use of TLS. Such regulatory aspects of the DNA damage tolerance could not have been observed with previous strategies using plasmid or randomly distributed DNA lesions, which shows the advantage of our new method. The very robust and precise integration of any modified DNA at any chosen locus of the yeast genome that we describe here is a powerful tool that will enable the exploration of many biological processes related to replication and repair of modified DNA.

## INTRODUCTION

Various exogenous and endogenous agents pose a constant threat to the genome of all organisms, resulting in DNA modifications such as abasic sites, DNA adducts (1), DNA crosslinks (intra-or inter-strand), DNA-protein crosslinks (2), presence of ribonucleotides (3). Unrepaired, these modifications present a serious challenge to a cell, as they may impair replication or give rise to deleterious mutations. In response to those threats, organisms have evolved many different mechanisms to deal with DNA damage.

Numerous repair systems exist that remove various modifications from DNA in an error-free manner (4). However, despite their efficient action, it is inevitable that some damages will persist long enough to be present during replication, which can lead to replication defects (replication blocks or delays, fork collapse) or alter replication fidelity. Therefore, to complete replication and maintain cell survival in the presence of residual DNA damages, cells have evolved two DNA Damage Tolerance (DDT) mechanisms: i) Translesion Synthesis (TLS), employing specialized DNA polymerases able to insert a nucleotide directly opposite the lesion. This pathway is potentially mutagenic due to the miscoding nature of most damaged nucleotides and to the low fidelity of the TLS polymerases (reviewed in (5)); ii) Damage Avoidance (DA, also named strand switch, copy choice or homology directed gap repair), a generally error-free pathway where the cells use the information of the sister chromatid to circumvent the lesion in an error-free manner (reviewed in (6)). The balance between error-free and error-prone mechanisms is important since it defines the level of mutagenesis during lesion bypass.

Decades of studies of DNA damage tolerance have yielded significant advances in our knowledge. It is well established that in eukaryotic cells, error-prone TLS is controlled by PCNA mono-ubiquitination while error-free DA is triggered by PCNA poly-ubiquitination. However, several questions remain regarding how the balance between these two pathways is controlled.

Over the years, many assays have been developed to study TLS and DA. Yet, their main limitation is that they cannot be used to monitor both TLS and DA at the same time. Genome-wide assays involving treatment with DNA damaging agents can be used to monitor toxicity and mutagenesis, but are blind to error-free events. The introduction of single DNA lesions onto replicative plasmids have been successfully used to monitor error-free and error-prone TLS (7–12). However, plasmid-based assays are not suited for the analysis of DA events, as during plasmid replication, when a lesion is encountered, replication fork uncoupling leads to full separation of the daughter strands in plasmids, while DA events require close proximity of the two sister chromatids (13).

To overcome the limitations of these approaches, we designed an assay to follow the fate of a single replication-blocking lesion introduced in the genome of a living cell. Our group has previously developed such assay in *Escherichia coli* (13, 14), and showed that indeed, such approach allows monitoring both TLS and DA events, and the interplay between these two tolerance mechanisms. It appeared necessary to develop a similar approach in eukaryotic cells in order to explore DNA damage tolerance in this kingdom of life. We chose to use the yeast *Saccharomyces cerevisiae* which provides an invaluable model due to the ease of genetic manipulation and high homology with several human genes. Furthermore, recent progress in construction of yeast strains with humanized genes and pathways opens up many possibilities for the study of human genes and processes in a simpler organismal context (15).

The method described here involves the site-specific integration of a vector containing a single DNA modification and a strand marker designed to distinguish the modified from the non-modified strand upon replication. A simple colorimetric assay is employed to monitor TLS and DA events.

We have used our method to insert two different UV lesions into the genome of the yeast *S. cerevisiae*. We confirm the involvement of several specialized DNA polymerases that has previously been observed using randomly distributed DNA lesions and plasmid assays. In addition, we show that impairing the DA pathway either at the control level (*ubc13*) or at the effector level (*rad51*), leads to an increase in the use of both error-free and mutagenic TLS. Such interplay between TLS and DA has never been observed before as it can only be evidenced on the chromosomal DNA and at the level of a single lesion. It shows the advantage of our method over currents approaches relying on plasmid-based assays or DNA lesions randomly distributed over the genome.

## MATERIAL AND METHODS

### Plasmids

pKM34 expresses the Cre integrase under the control of a tetracycline-repressible TetO-*CYC1* promoter and carries *TRP1* selectable marker. The vector is derived from pCM185 (16) by cloning Cre recombinase from pSH68 (17) into the HpaI-PstI restriction sites.

pLL43 and pKM71 are derived from pUC19 plasmid and contain: the ampicillin resistance gene and pMB1 bacterial replication origin from the pUC19, the yeast *LEU2* marker from pSH68, and the 5’ end of *lacZ* gene cloned from *E. coli* MG1655 strain in fusion with the *lox71* site under control of *pTEF* promoter from pYM-N18 (18). pRS413 (19) plasmid, carrying *HIS3* selectable marker, serves as an internal control for transformation efficiency.

### Construction of vectors carrying a single lesion

Duplex plasmids carrying a single lesion were constructed following the gap-duplex method previously described (20). An oligonucleotide (5’-GCAAGTTAACACG) containing no lesion, a thymine-thymine pyrimidine(6–4) pyrimidone photoproduct [TT(6-4)], or a cyclobutane pyrimidine dimer (TT-CPD) lesion (underlined) was inserted into the gapped-duplex pLL43/pKM71 creating an in frame *lacZ* 5’ gene fragment.

### Strains

All strains used in the present study are derivative of strain EMY74.7 (21) (MATa *his3-Δ1 leu2-3,112 trp1-Δ ura3-Δ met25-Δ phr1-Δ rad14-Δ msh2Δ∷hisG).* Gene disruptions were achieved using PCR-mediated seamless gene deletion (22), URAblaster (23), or delitto perfetto (24) techniques. All strains used in the study are listed in Table S1.

All strains carry the plasmid pKM34 expressing Cre recombinase, and the chromosomal integration cassette (*lox66-3’lacZ-MET25*) containing the 3’ end of *lacZ* gene cloned from *E. coli* MG1655 strain in fusion with the *lox66* site. Integration cassettes were placed in two alternative locations within the yeast genome: near *ARS306* or *ARS606*, using *MET25* as selection marker.

### Yeast Transformation

Plasmids carrying UV lesions (or control plasmids without lesion) were introduced into yeast cells by electroporation. Yeast cells were grown overnight to stationary phase in synthetic defined yeast medium (yeast nitrogen base, ammonium sulfate, glucose) lacking tryptophane (SD-TRP) with 5 μg/ml doxycycline. The overnight culture was inoculated into 100 ml of yeast extract/peptone/dextrose medium (YPD) to reach OD_600_=0.3. The cells were grown until OD _600_ =1.6 and harvested by centrifugation. The pellet was washed twice with 50 ml of cold water and once with 50 ml of cold electroporation buffer (1 M sorbitol/1 mM CaCl_2_). Cells were then incubated for 30 min at 30°C in conditioning buffer (0.1 M LiAc/10 mM DTT), collected by centrifugation, washed one more time with 50 ml of electroporation buffer, and then resuspended in 100–200 μl of the same buffer to reach 1 ml volume.

DNA mix (100 ng of integrative plasmid with/without lesion, 100 ng of transformation control plasmid pRS413, and 12 μg of denatured carrier DNA) was added to 400 μl of cell suspension. Cells were electroporated at 2.5 kV/25 mF/ 400 Ω in a BioRad GenePulser using 0.2 cm electrode gap cuvette. Typical electroporation time constant ranged from 3.0 to 4.5 ms. After electroporation, cells were suspended in 6 ml of 1:1 mix of 1 M sorbitol: YPDplus medium (Zymo Research) and incubated at 30°C for 1 h. Finally, the cells were collected, resuspended in 5 ml 1M sorbitol, and plated on selective media using the automatic serial diluter and plater EasySpiral Dilute (Interscience).

Part of the cells were plated on synthetic defined yeast medium (yeast nitrogen base, ammonium sulfate, glucose, agar) lacking histidine (SDa-HIS) to measure the transformation efficiency of plasmid pRS413 (internal transformation standard), and the rest was plated on synthetic defined yeast medium lacking leucine (SDa-LEU) (with 80 μg/ml X-gal and 10 μg/ml doxycycline) to select for integrants and visualize TLS and DA events as sectored blue-white and white colonies, respectively. One should note that the lesion is not repaired in the recipient strains. Therefore, after the first round of replication, whether the lesion has been bypassed by TLS or DA, one of the daughter cells will inherit the lesion that will have to be bypassed again during the second cycle of replication. A lesion bypassed by TLS at the second round of replication will lead to a blue colony and will be scored as TLS. We expect the mother cell where the lesion remains to be rapidly diluted over time. Colonies were counted using the Scan 1200 automatic colony counter (Interscience). On average, 2000-5000 colonies were counted per experiment. Lesion tolerance rates were calculated as the relative integration efficiencies of damaged vs non-damaged vectors normalized by the transformation efficiency of pRS413 plasmid in the same experiment. DA events are estimated by subtracting TLS events from the total lesion tolerance events. The integration efficiency is about 100 CFU/ng of non-damaged vector (3×10^4^ CFU/pmol) for our parental strain *(msh2 rad14),* which corresponds to 0.02% of electroporated cells. Each experiment was performed in at least 3 independent replicates, carried out on different days with different batches of competent cells.

A point by point protocol is provided in the supplementary material.

## RESULTS

### Site-specific integration into the yeast genome

In order to overcome the limitations previously described, we developed an assay that allows one to follow the fate of a single replication-blocking lesion in the yeast genome. This technique is based on a non-replicative plasmid containing a single lesion, which is stably integrated into one of the chromosomes using site-specific recombination, as previously described for *E. coli* (13, 14) (Fig.1). After testing several integration strategies (see supplementary information and supplementary table S3), we chose a modified version of the Cre/*lox* system involving Leu-Element/Right-Element (LE/RE) *lox* site mutants (Fig.1C). Recombination between LE (*lox61*) and RE (*lox71*) *lox* mutants produces a wild-type *loxP* site as well as a LE+RE double mutant *lox* site that is not recognized by Cre (25), thus preventing excision of the plasmid once integrated into the chromosome. In addition, if several plasmids enter the cell, once one of them is integrated, the remaining ones cannot be exchanged on the chromosome. Additionally, we placed the Cre recombinase under the control of the doxycycline repressible promoter (Tet-off) so it can be turned off after integration has occurred.

Following ectopic expression of Cre recombinase (pKM34), the plasmid carrying a lesion is introduced by electroporation into a recipient *S. cerevisiae* strain containing a chromosomal integration cassette. The plasmid contains a selectable marker (*LEU2*), and a single lesion located within the 5-end of the *lacZ* gene fused to a *lox71* site. The chromosomal integration site contains the 3-end of *lacZ* fused to *lox66* site, so that following the precise integration a full-length functional ß-galactosidase gene (*lacZ*) is restored (Fig.1).

We placed the chromosomal integration site close to an early replication origin in two different orientations so we can choose to introduce the lesion either on the leading or the lagging strand (Fig.1C and Fig.S1). To rule out any possible bias due to the choice of the integration site, we chose two different integration sites, on chromosomes III and VI (close to the early replication origins *ARS306* or *ARS606*) (Fig. 1B). The non-damaged strand contains a +2 frameshiu inactivating the *lacZ* gene, serving as a genetic marker for strand discrimination (Fig. 1C). After electroporation of the vector, cells are plated on selective indicator plates (SDa-LEU, X-gal) before the first cell division occurs. The lesion is placed in such sequence context, that all in-frame TLS events, both error-free and mutagenic, result in a functional *lacZ* gene (blue colony sectors), while DA events result in inactivated *lacZ* gene (white colony) (Fig.1D). Integration of lesion free vectors gives rise to essentially only sectored colonies (Fig.1D).

**Figure 1:**
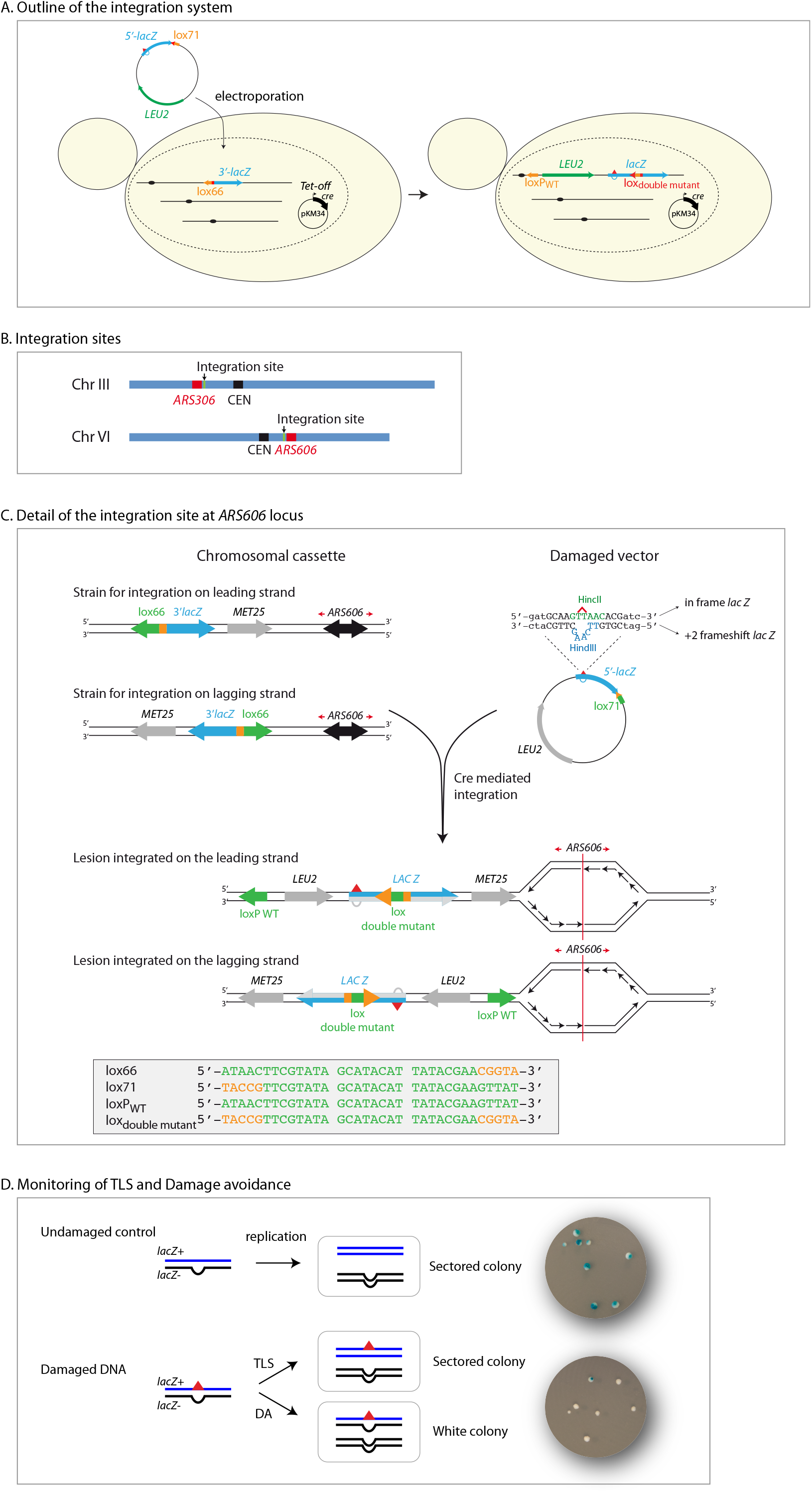
Introduction of a single lesion into a defined location in the yeast genome. A. Outline of the integration system: A non-replicative plasmid containing a single lesion is integrated into one of the yeast chromosomes (III or VI) using Cre/*lox* site-specific recombination. The integrative vector carrying a selection marker (*LEU2*) and the 5’-end of the *lacZ* reporter gene containing a single lesion is introduced into a specific locus of the chromosome with the 3’-end of the *lacZ*. The precise integration of the plasmid DNA into the chromosome restores a functional *lacZ* gene, enabling the phenotypical detection of TLS and DA events (as blue and white colonies on X-gal indicator media). Plasmid pKM34 expresses the Cre recombinase under the control of the doxycycline repressible promoter (Tet-off). B. Integration sites: we have designed two different integration sites, one on chromosome III in the vicinity of *ARS306*, one on chromosome VI in the vicinity of *ARS606*. C. Detail of the integration site at *ARS606*: The recipient strain carries a chromosomal cassette, containing 3’-end of the *lacZ* gene fused to a *lox66* site. The cassette is placed near a robust, early firing origin of replication (*ARS606*) in one of two possible orientations, which permits after vector integration to monitor the replication of the locus containing a single lesion as a leading or lagging strand template. The non-replicative vector contains a selection maker (*LEU2*) and the 5’-end of the *lacZ* reporter gene containing a single lesion (red triangle), fused to a *lox71* site. The non-damaged opposite strand contains a + frameshiu inactivating the *lacZ* gene, serving as a genetic marker to enable strand discrimination. The lesion is placed in such a sequence context, that an error-free TLS events results in HincII restriction site, while the non-damaged strand opposite the lesion contains a HindIII restriction site. To prevent excision of the lesion containing region after integration, leu element/right element (LE/RE) *lox* site mutants have been used. Recombination between LE (*lox61*) and RE (*lox71*) lox mutants produces a wild-type *loxP* site as well as a LE+RE double mutant *lox* site within the *lacZ* gene that is not recognized by Cre. WT element of lox site are show in green, while mutated elements are shown in orange. The detailed sequence of the lox sites is shown in the grey box. Detail of integration site at *ARS306* is shown in Fig. S1. D. Monitoring TLS and Damage avoidance: chromosomal integration of undamaged of *lac*-/*lac*+ heteroduplex constructs lead to sectored colonies on indicating media. Replication of the damaged heteroduplex yields a *lac*+ event when the lesion is bypassed by TLS, whereas complementary strand replication yields a *lac* – event. Damage Avoidance (DA) events lead to two lac-events and therefore to the formation of white colonies.

PCR analysis and sequencing confirmed that all colonies obtained on selective plates result from precise integration of the vector into the chromosomal integration site. No colonies were observed after transformation of a strain not expressing Cre recombinase or without chromosomal integration site.

### Bypass of UV lesions by Translesion Synthesis

To validate our system, we constructed 3 integration vectors, containing no lesion, a TT-CPD lesion (cyclobutane pyrimidine dimer) and a TT(6-4) lesion (thymine-thymine pyrimidine(6-4)pyrimidone photoproduct). In order to focus on lesion tolerance mechanisms, we inactivated the repair mechanisms in our parental strain (namely nucleotide excision repair: *rad14*, and photolyase: *phr1*), as well as the mismatch repair system (*msh2*), to avoid the repair of the strand marker. Tolerance events are calculated as the ratio of colonies resulting from the integration of the damaged vector versus the lesion-free one.

We integrated the constructs containing a single CPD or TT(6-4) lesion as well as a construct with no lesion. The data express the level of tolerance events (either TLS or DA) relative to the non-damaged construct. After integration of the constructs containing a single CPD or TT(6-4) lesion, no reduction of survival was observed compared to the lesion-free construct (Fig.2). Integration of the heteroduplex containing a single CPD lesion leads to 55% of sectored blue colonies representing TLS events. For TT(6-4) lesion, 4% of TLS events were observed. Those results are in agreement with a previous report by Gibbs et al. (9), where the authors used gapped-circular vectors containing a single lesion within a short single-stranded region. For the two designed chromosomal integration sites, we introduced the two lesions in the leading strand and in the lagging strand (relative to the closest replication origin, *ARS306* and *ARS606*). As shown on Fig. 2, we observed no differences between the leading and lagging strand nor between the two integration sites. Since no difference was observed between *ARS306* and *ARS606* nor between leading and lagging strand integrations, for the next graphs (Fig. 3 and 4) we have shown the average of data observed at the two integration sites and in the two orientations.

**Figure 2:**
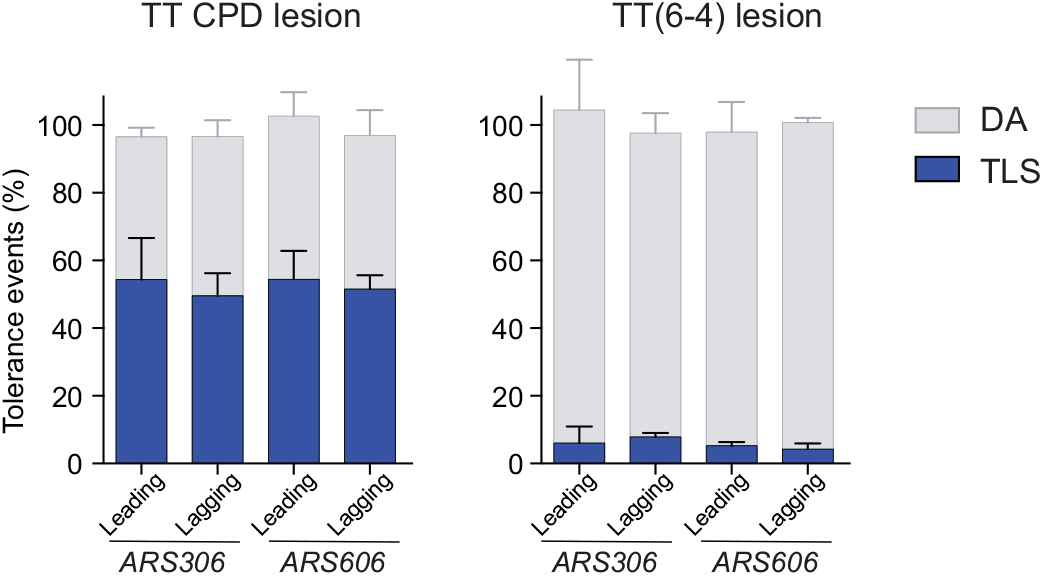
Partitioning of DDT pathways at different integration sites: The graph represents the partion of DDT pathways for two UV lesions, in our parental strains deficient in repair mechanisms (*phr1Δ rad14Δ msh2Δ∷hisG*), where the lesion is introduced at two different integration sites (close to *ARS306* or *ARS606*) either on the leading or the lagging strand. Tolerance events represent the percentage of cells able to survive in presence of the integrated lesion **compared to the lesion-free control**. The data represent the average and standard deviation of at least three independent experiments. Unpaired *t*-test was performed to compare TLS and DA values from the different integration sites: no significant difference was observed (p > 0.2).

**Figure 3:**
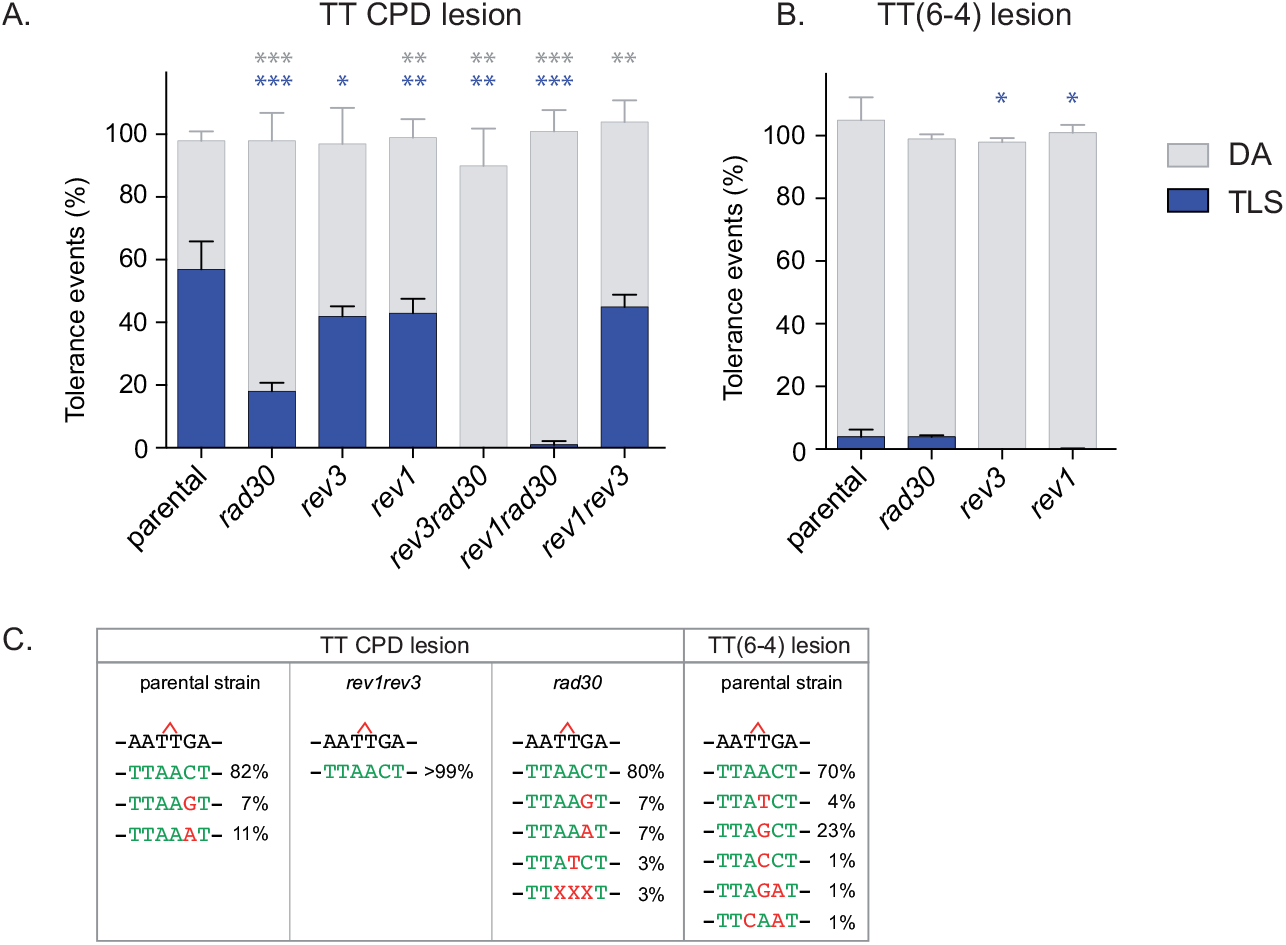
Partitioning of DDT pathways in a strain deficient in TLS polymerases. The graph represents the partion of DDT pathways for two UV lesions, in strains deficient in TLS polymerases (Rev1, Pol η/*rad30*, Pol ζ/*rev3*). Tolerance events represent the percentage of cells able to survive in presence of the integrated lesion **compared to the lesion-free control**. The data represent the average and standard deviation of at least three independent experiments in which the lesion has been inserted either in the leading or the lagging strand. Unpaired *t*-test was performed to compare TLS and DA values from the different mutants to the parental strain. \**P* < 0.05; \*\**P* < 0.005; \*\*\**P* < 0.0005. All strains are *phr1Δ rad14Δ msh2Δ∷hisG.* A. In a parental strain over 50% of events observed across a CPD lesion are TLS events. DNA polymerase is responsible for the majority of CPD lesion bypass. However, in its absence TLS bypass of this lesion is still possible through the combined action of polymerase and Rev1. In the absence of pol, removal of either Rev1 or pol ζ completely abolishes TLS. B. For the TT(6-4) lesion, DA is the major tolerance pathway. Majority of TLS events through TT(6-4) lesion depend on pol and Rev1, while pol rarely contributes to the bypass of this lesion. C. Nucleotides inserted opposite the CPD and TT(6-4) lesions in the different mutants of Polymerases. Error-free and mutagenic insertions are represented in green and red respectively. A more detailed mutagenesis spectrum is provided in supplementary Table S2.

**Figure 4:**
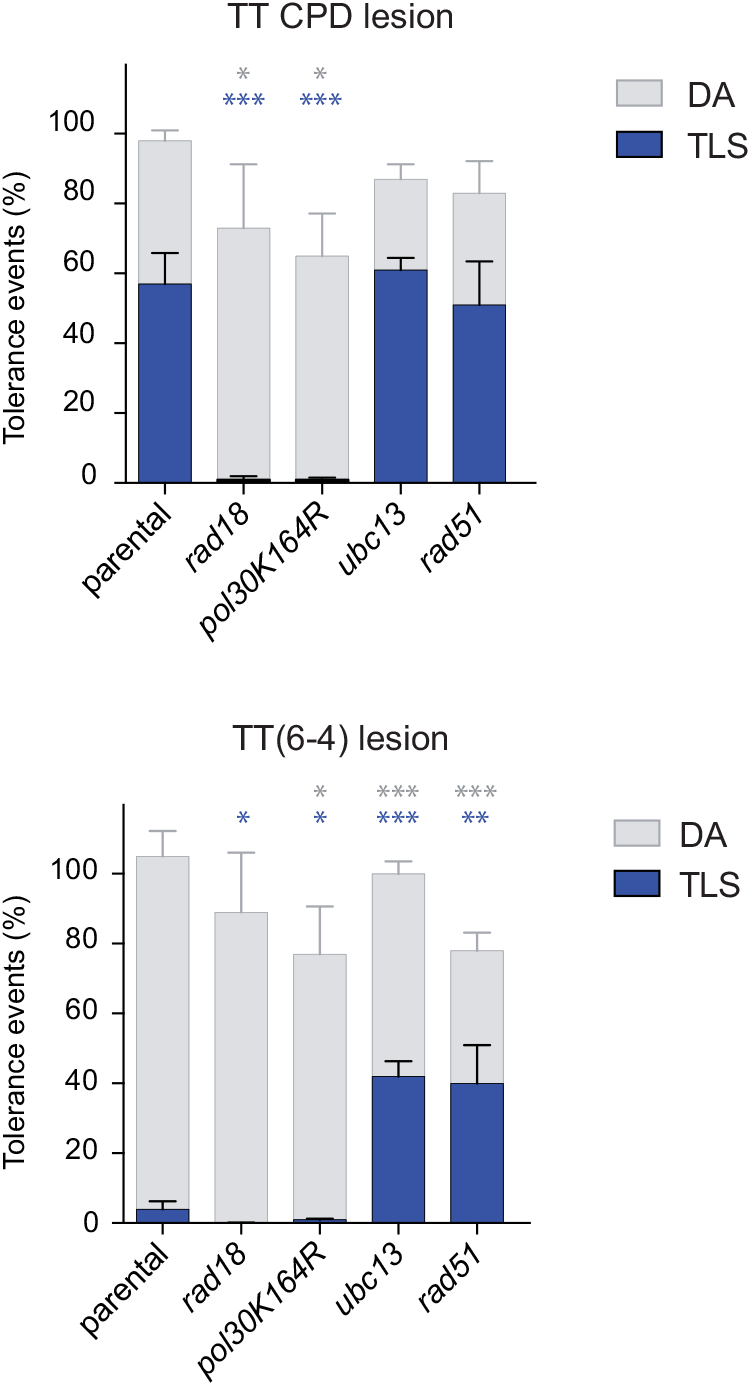
Partitioning of DDT pathways in the absence of PCNA ubiquitylation. Bypass of TT-CPD (A) and TT(6–4) (B) lesion in strains deficient in PCNA ubiquitylation (*rad18* or *pol30-KtiR*) or polyubiquitylation (*ubc13*). Tolerance events represent the percentage of cells able to survive in presence of the integrated lesion compared to the lesion-free control. The data represent the average and standard deviation of at least three independent experiments in which the lesion has been inserted either in the leading or the lagging strand. Unpaired *t*-test was performed to compare TLS and DA values from the different mutants to the parental strain. \**P* < 0.05; \*\**P* < 0.005; \*\*\**P* < 0.0005. All strains are *phr1Δ rad14Δ msh2Δ∷hisG.* In the absence of PCNA ubiquitylation (*rad18, pol30 KtiR*) we observed a decrease in TLS. The remaining low level of TLS is probably due to polymerase interactions with PCNA ring not involving ubiquitin moiety. In the absence of PCNA polyubiquitylation (*ubc13*), a small increase of the TLS bypass of the CPD lesion is observed, while Pol-mediated bypass of the TT(6-4) lesion increased more than 10 fold. The absence of recombinase Rad51 results in a similar increase in TLS.

### Modulation of TLS by the specialized DNA polymerases

In the absence of Pol (*rad30*), TLS at CPD lesion is strongly reduced to ~18% (Fig. 3). The remaining TLS in the absence of Pol is dependent on *REV1* and *REV3* as inactivation of either of these genes in combination with *rad30* leads to an almost complete suppression of TLS events. Despite the drop of 63% in the rate of TLS at the CPD in the absence of Pol, we observe no loss of survival. This is consistent with the study by Abdulovic and Jinks-Robertson (26) where the authors demonstrated that *rad30* strain is not sensitive to low UV doses. We can therefore conclude that in the presence of low levels of DNA damage homologous recombination-dependent mechanism (DA) can fully compensate for the absence of specialized polymerase.

In the presence of Pol, inactivation of either *REV1* or *REV3* leads to a milder reduction of TLS at a CPD lesion. Both genes are epistatic as the inactivation of both *rev3* and *rev1* leads to the same decrease of TLS. It is interesting to note that Pol - m?ediated TLS and Rev1-Rev3-mediated TLS are independent from each other and seem compartmentalized: the drop in TLS in the absence of Pol cannot be compensated by Rev1-Rev3 TLS and vice versa.

*RAD30* inactivation has no effect on TLS at TT(6-4) lesions. However, *REV1* or *REV3* (or both) inactivation leads to a complete suppression of TLS at this lesion, showing again the epistasis of both genes in the bypass of this lesion.

Molecular analysis of colonies obtained after lesion integration (Fig. 3C and Table S2) shows that insertion opposite the CPD lesion is 100% error-free in the presence of Pol, as expected from the specificity of this polymerase for this lesion (27). However, it is interesting to note that even in the presence of Pol, the bypass of CPD is mutagenic in 18% of the cases due to mis-elongation by Rev1-Pol. The insertion step becomes mutagenic (5%) in the absence of Pol. Overall mutagenesis (both at the insertion and elongation steps) is almost completely abolished in the absence of *rev3* and *rev1*. The bypass of the TT(6-4) lesion is mutagenic in 30% of the case, mostly due to misincorporation at the insertion step.

Altogether, these results confirm the involvement of TLS polymerases in the bypass of UV lesions that was previously obtained with replicative or gapped plasmids (9, 12). They show that our method by which we introduce a single lesion in the genome is suitable to study TLS. In addition, we show that a decrease in TLS caused by the absence of one (or several) specialized DNA polymerase(s) is fully compensated by a concomitant increase in the DA process, avoiding any decrease in the cell survival.

### PCNA mono-ubiquitination is required for TLS

It is known that the balance between TLS and DA is regulated by post-translational modifications of PCNA that occur in response to DNA damage. PCNA mono-ubiquitination (at Lysine 164) is mediated by Rad6/Rad18 and favors TLS (28). PCNA poly-ubiquitination depends on Mms2-Ubc13 ubiquitin-conjugating complex and the ubiquitine-ligase activity of Rad5, and is important for DA (29). Since our system was designed to explore the balance between error-prone and error-free lesion bypass pathways, we investigated the role of PCNA ubiquitination on the bypass of our UV lesions. We introduced our two UV lesions in strains were PCNA cannot be ubiquitinated, either by the inactivation of *RAD18*, or by the mutation of Lysine 164 of PCNA (*pol30*-K164R) (Fig. 4). In both situations, in the absence of PCNA ubiquitination, the TLS level at CPD and TT(6-4) lesions is almost completely abolished. It appears therefore that PCNA ubiquitination is an absolute requirement for TLS.

Interestingly, in the absence of any DNA-damaging treatment, the presence of a single replication-blocking lesion is sufficient to generate the signal required to trigger Rad6-Rad18-mediated PCNA ubiquitination which is clearly necessary to promote TLS.

It is also interesting to note that DA is still possible in the absence of PCNA ubiquitination since we still observe a great proportion of white colonies in the *rad18* and the *pol30-K164R* mutant. However, while the drop in TLS caused by the absence of one or multiple DNA polymerases was fully compensated by DA mechanisms (Fig. 3 and previous paragraph), the drop of TLS induced by the lack of PCNA ubiquitination can only be partially compensated by DA, leading to a reduced survival (Fig. 4). This shows that DA partially depends on PCNA ubiquitination.

### Lack of PCNA poly-ubiquitination favors TLS

We then looked at how PCNA poly-ubiquitination could affect the ratio DA/TLS (Fig. 4). In the absence of PCNA poly-ubiquitination (*ubc13* strain), we observed no significant effect on the bypass of CPD lesion. On the other hand, we observed a more than 10 fold increase in the Pol - mediated TLS at the TT(6-4) lesion, reaching more than 40% (Fig. 4). In the absence of PCNA poly-ubiquitination, DA is reduced and is compensated by an increase in TLS. Such phenomenon has previously been observed in *E. coli* where we showed that a defect in homologous recombination led to increased TLS (30) using a similar approach. Only the monitoring of a single DNA lesion inserted in the genomic DNA is able to reveal such interplay between DA and TLS.

To confirm that the increase in TLS was indeed due to a decrease in DA, we inactivated the Rad51 recombinase which is a key actor in the DA mechanism (31–33). In the *rad51* strain, we observed the same 10-fold increase in TLS rate at the TT(6-4) lesion (Fig. 4). Such effect of *rad51* deletion has previously been observed by Morrison and Hastings (34). This confirms that affecting the DA process, either at its regulation level (*ubc13*) or at its effector level (*rad51*) leads to an increase in TLS.

As we did not observe a similar increase in TLS at the CPD lesion, we hypothesized that the competion between DA and TLS occurs behind the fork, during a gap filling reaction. Following the encounter with a blocking lesion, a repriming event generates a single-strand DNA gap that will be filled post-replicatively (35, 36, 37). The majority of CPD lesions is efficiently bypassed by Pol at the fork. Only the small fraction that is bypassed by Pol behind the fork, is in competion with *UBC13*-dependant DA mechanisms for the gap-filling reaction. This hypothesis will need to be confirmed by the use of other strongly-blocking DNA lesions that are bypassed by Pol. The difference we observe between CPD and TT(6-4) is in agreement with the work of Fasullo et al. (38) where they show that *UBC13* is not required for UV-associated sister chromatid exchange (SCE), but is required in response to MMS or 4-NQO treatment. When UV-irradiating the cells, they generate a majority of CPD lesions and therefore do not see the requirement of *UBC13* at this lesion. Our approach confirms that *UBC13* has no effect on the bypass of CPD lesions, but shows in addition that *UBC13* affects the bypass of TT(6-4) lesions. This shows the higher resolution of our method compared to global damage generated by UV irradiation or other treatments.

In the *ubc13* or *rad51* strains, some DA still persists as we still observe a significant number of white colonies. These colonies could arise from *RAD51*-independent template switching mechanisms that rely on *RAD52* or *RAD5* (39, 40). Another possibility is the occurrence of damaged chromatid loss as previously observed in *E. coli* (41). By adding markers on the damaged and non-damaged strand, we will be able to explore and characterize these DA events as previously achieved in *E. coli* (41).

## DISCUSSION

Our goal was to develop a method to explore the mechanisms employed by living cells to deal with DNA alterations. Over the years, many assays have been developed to study error-prone TLS or error-free DA. However, their main limitation is that they cannot be used to monitor both TLS and DA simultaneously. Assays measuring chromosomal mutagenesis after treatment with mutagenic agents are blind to error-free events. Plasmid-based systems have been successfully used to monitor error-free and error-prone TLS (7–12). They are, however, not suited for the analysis of DA events (13).

In the present paper, we describe a method that overcomes these limitations by making it possible to analyze the fate of a single DNA modification inserted in the genome of a yeast cell. We have used this method to introduce a single UV lesion (TT(6-4) or CPD) into the genome of *S. cerevisiae* and studied its bypass by DNA damage tolerance pathways. Several factors have been proposed to regulate the interplay between TLS and DA, among them the nature of the lesion and the post-translational modification of PCNA. However, no high-resolution assays were able to monitor both TLS and DA simultaneously at the level of a single lesion in the genome. Using our method, we were able to show that the proportion of TLS *vs.* DA is dependent on the lesion: while TLS represents ~55% of the tolerance pathways for CPD, it represents only ~4% for TT(6-4). This difference is due to the presence of Pol and the specificity of its more open active site that can accommodate CPD lesions (42). For both lesions, no toxicity is observed and DA complements TLS pathway in order to recover 100% of survival (as compared to the integration of the lesion-free control vector). We showed that in the absence of TLS polymerases, DA mechanisms could fully compensate for the lack of TLS avoiding any drop in survival. However, when PCNA ubiquitination was abolished, TLS was almost completely suppressed but could only be partially compensated by DA, showing that DA partially depends on PCNA ubiquitination.

More interestingly, we showed that a defect in the DA pathway leads to an increase in TLS. Indeed, at the TT(6-4) lesion, when the DA pathway is affected either by the inactivation of *ubc13* or of *rad51*, it is compensated by a 10 fold increase in TLS. This increase of TLS due to a defect in DA can only be revealed by our method. Previously used approaches based on randomly distributed lesion (e.g. UV irradiation) could reveal an increase in mutagenesis, but were blind to error-free processes (including DA and error-free TLS). It has previously been reported that *ubc13* inactivation led to a ~2-fold increase in UV-induced mutagenesis (43), reflecting the low fraction of mutagenic TLS events. We report here a >10-fold increase in the use of Pol-mediated TLS in the *ubc13* strain, as our system allows measuring both mutagenic and error-free TLS events. Plasmid-based assays designed to monitor error-free TLS have been used, but they are inappropriate substrates to monitor DA: due to their limited size, the full unwinding of the two DNA strands prevents homologous recombination with the sister chromatid as previously evidenced in *E. coli* (13). Indeed, the inactivation of *ubc13* does not lead to any increase of TLS at a single lesion bypassed on a plasmid system in *S. cerevisiae* (44).

The increase of TLS in response to inhibition of DA evidenced here could not be observed before in yeast since no assay was able to simultaneously monitor TLS and DA, and therefore the interplay between these two mechanisms. The method described here has the potential to unveil several new aspects of the DNA damage tolerance. Many factors may affect the DNA damage tolerance pathway choice, including lesion type, sequence context, location in the genome, chromatin state, cell cycle stage, and transcriptional activity. The versatility of our assay will facilitate the exploration of the impact of those factors. Our assay allows inserting any type of DNA lesion or modification at any desired location in the yeast genome. By placing the DNA damage in centromeric or telomeric regions, highly/poorly transcribed regions, hetero/eu-chromatin regions, near fragile sites, it will be possible to determine how these positions affect the balance between error-free and mutagenic lesion bypass. In addition, the integration site can be placed in two orientations relative to the closest replication origin, so that the lesion can be placed on the leading or lagging strand. Similarly, we can choose to insert the lesion on the transcribed, or on the non-transcribed strand.

This method opens the way to exploration of lesion bypass and single-strand gap repair in the same manner engineering nucleases such as HO or I-SceI has boosted the exploration of double strand breaks repair (45).

This method is not limited to DNA Damage tolerance, but can also be used to explore several mechanisms related to DNA maintenance, such as repair of DNA adducts, repair of DNA crosslinks, effect of ribonucleotides inserted into the DNA (3), or effect of DNA-protein crosslinks (2). The possibility to integrate vectors carrying any type of DNA damage or chemical group broadens the applicability of our approach beyond the field of DNA damage repair and tolerance. Being able to locate a single modification at a specific locus of the genome will enable powerful molecular analysis at the resolution of a single replication fork.

