## Supplementary information for "iDamage: a method to integrate modified DNA into the yeast genome"

### Figure S1

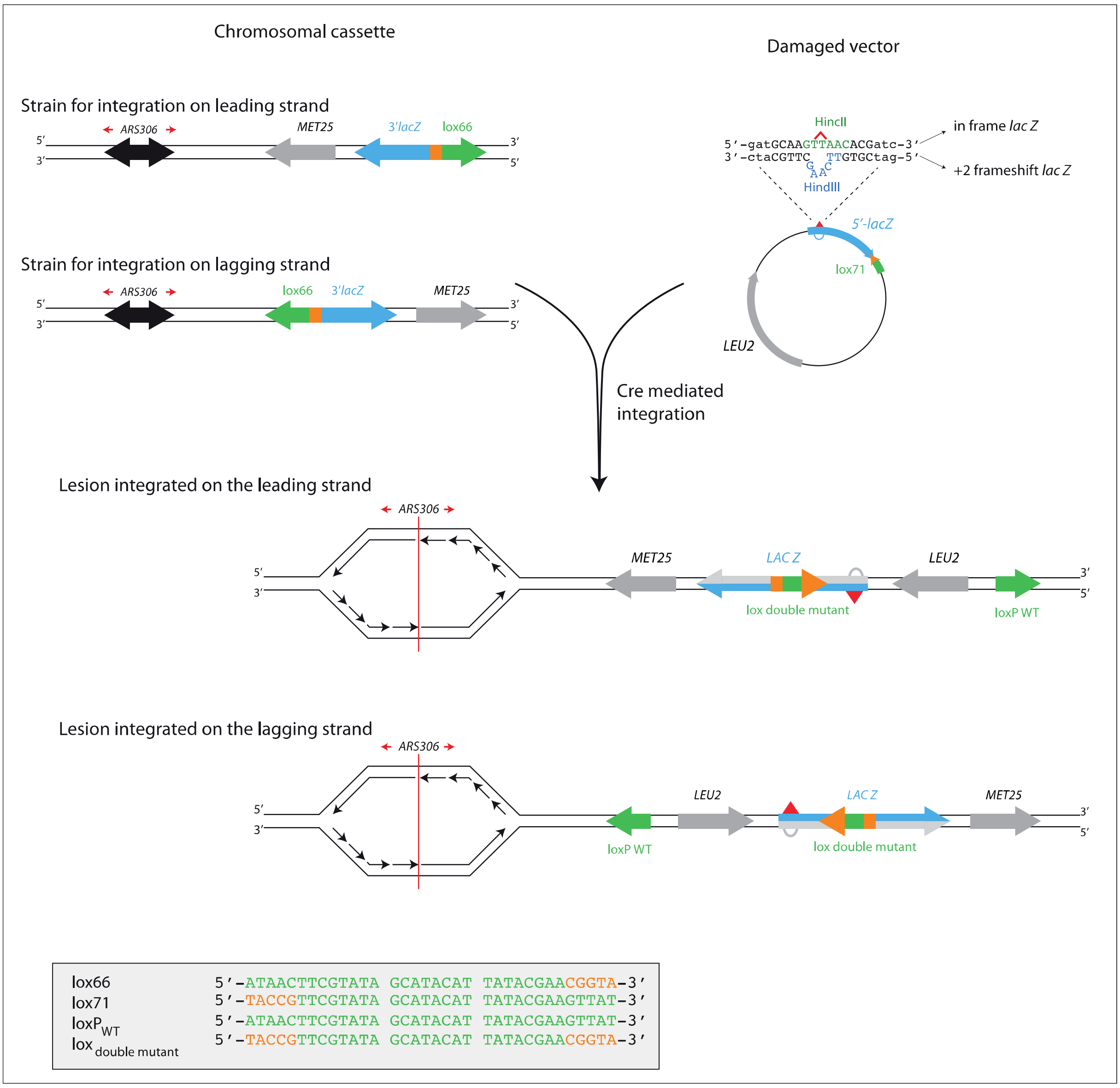


Detail of the integration site at *ARS306:* The recipient strain carries a chromosomal cassette, containing 3'-end of the *lacZ* gene fused to a *lox66* site. The cassette is placed near a robust, early firing origin of replication (*ARS306*) in one of two possible orientations, which permits after vector integration to monitor the replication of the locus containing a single lesion as a leading or lagging strand template.

The non-replicative vector contains a selection maker (*LEU2*) and the 5'-end of the *lacZ* reporter gene containing a single lesion (red triangle), fused to a *lox71* site. The non-damaged opposite strand contains a +2 frameshift inactivating the *lacZ* gene, serving as a genetic marker to allow strand discrimination. The lesion is placed in such a sequence context, that an error-free TLS events results in HincII restriction site, while the non-damaged strand opposite the lesion contains a HindIII restriction site.

To prevent excision of the lesion containing region after integration, left element/right element (LE/RE) *lox* site mutants have been used. Recombination between LE (*lox61*) and RE (*lox71*) lox mutants produces a wild-type *loxP* site as well as a LE+RE double mutant *lox* site within the *lacZ* gene that is not recognized by Cre. WT element of lox site are show in green, while mutated elements are shown in orange. The detailed sequence of the lox sites is shown in the grey box.

**Table S1: Strains used in the study**

| Strain | Relevant genotype<br>(All strains are: *MATa his3-Δ1 leu2-3,112 trp1-Δ ura3-Δ met25-Δ rad14-Δ phr1-Δ msh2Δ::hisG*) |
|---|---|
| SC49 | III(75494-75499)::(*lox66-3'lacZ-MET25*/lead) |
| SC51 | III(75494-75499)::(*lox66-3'lacZ-MET25*/lag) |
| SC53 | VI(167260-167265)::(*lox66-3'lacZ-MET25*/lag) |
| SC55 | VI(167260-167265)::(*lox66-3'lacZ-MET25*/lead) |
| SC82 | *rev1-Δ* VI(167260-167265)::( *lox66-3'lacZ-MET25*/lag) |
| SC83 | *rev1-Δ* VI(167260-167265)::(*lox66-3'lacZ-MET25*/lead) |
| SC84 | *rev1-Δ* III(75494-75499)::(*lox66-3'lacZ-MET25*/lag) |
| SC85 | *rev1-Δ* III(75494-75499)::(*lox66-3'lacZ-MET25*/lead) |
| SC86 | *rad30Δ::hisG* VI(167260-167265)::(*lox66-3'lacZ-MET25*/lag) |
| SC87 | *rad30Δ::hisG* VI(167260-167265)::(*lox66-3'lacZ-MET25*/lead) |
| SC88 | *rad30Δ::hisG* III(75494-75499)::(*lox66-3'lacZ-MET25*/lag) |
| SC89 | *rad30Δ::hisG* III(75494-75499)::(*lox66-3'lacZ-MET25*/lead) |
| SC181 | *rev3Δ::hisG* VI(167260-167265)::(*lox66-3'lacZ-MET25*/lag) |
| SC182 | *rev3Δ::hisG* VI(167260-167265)::(*lox66-3'lacZ-MET25*/lead) |
| SC183 | *rev3Δ::hisG* III(75494-75499)::(*lox66-3'lacZ-MET25*/lag) |
| SC184 | *rev3Δ::hisG* III(75494-75499)::(*lox66-3'lacZ-MET25*/lead) |
| SC171 | *rad30Δ::hisG* VI(167260-167265)::(*lox66-3'lacZ-MET25*/lag) |
| SC172 | *rad30Δ::hisG* VI(167260-167265)::(*lox66-3'lacZ-MET25*/lead) |
| SC173 | *rad30Δ::hisG* III(75494-75499)::(*lox66-3'lacZ-MET25*/lag) |
| SC174 | *rad30Δ::hisG* III(75494-75499)::(*lox66-3'lacZ-MET25*/lead) |
| SC163 | *rev1-Δ rad30Δ::hisG* VI(167260-167265)::(*lox66-3'lacZ-MET25*/lag) |
| SC164 | *rev1-Δ rad30Δ::hisG* VI(167260-167265)::(*lox66-3'lacZ-MET25*/lead) |
| SC165 | *rev1-Δ rad30Δ::hisG* III(75494-75499)::(*lox66-3'lacZ-MET25*/lag) |
| SC166 | *rev1-Δ rad30Δ::hisG* III(75494-75499)::(*lox66-3'lacZ-MET25*/lead) |
| SC159 | *rev1-Δ rev3Δ::hisG* VI(167260-167265)::(*lox66-3'lacZ-MET25*/lag) |
| SC160 | *rev1-Δ rev3Δ::hisG* VI(167260-167265)::(*lox66-3'lacZ-MET25*/lead) |
| SC161 | *rev1-Δ rev3Δ::hisG* III(75494-75499)::(*lox66-3'lacZ-MET25*/lag) |
| SC162 | *rev1-Δ rev3Δ::hisG* III(75494-75499)::(*lox66-3'lacZ-MET25*/lead) |
| SC151 | *ubc13-Δ* VI(167260-167265)::(*lox66-3'lacZ-MET25*/lag) |
| SC152 | *ubc13-Δ* VI(167260-167265)::(*lox66-3'lacZ-MET25*/lead) |
| SC153 | *ubc13-Δ* III(75494-75499)::(*lox66-3'lacZ-MET25/*lag) |
| SC154 | *ubc13-Δ* III(75494-75499)::(*lox66-3'lacZ-MET25*/lead) |
| SC203 | *rad18Δ::hisG* VI(167260-167265)::(*lox66-3'lacZ-MET25*/lag) |
| SC204 | *rad18Δ::hisG* VI(167260-167265)::(*lox66-3'lacZ-MET25*/lead) |
| SC205 | *rad18Δ::hisG* III(75494-75499)::(*lox66-3'lacZ-MET25*/lag) |
| SC206 | *rad18Δ::hisG* III(75494-75499)::(*lox66-3'lacZ-MET25*/lead) |
| SC236 | *pol30-K14R* VI(167260-167265)::(*lox66-3'lacZ-MET25*/lag) |
| SC237 | *pol30-K14R* VI(167260-167265)::(*lox66-3'lacZ-MET25*/lead) |
| SC238 | *pol30-K14R* III(75494-75499)::(*lox66-3'lacZ-MET25*/lag) |

| | |
|---|---|
| SC239 | *pol30-K14R* III(75494-75499)::(*lox66-3'lacZ-MET25*/lead) |
| SC254 | *rad51Δ::hyg* VI(167260-167265)::(*lox66-3'lacZ-MET25*/lag) |
| SC255 | *rad51Δ::hyg* VI(167260-167265)::(*lox66-3'lacZ-MET25*/lead) |
| SC256 | *rad51Δ::hyg* III(75494-75499)::(*lox66-3'lacZ-MET25/*lag) |
| SC257 | *rad51Δ::hyg* III(75494-75499)::(*lox66-3'lacZ-MET25*/lead) |
| SC348 | *ubc13-Δ rad30Δ::hisG* VI(167260-167265)::(*lox66-3'lacZ-MET25*/lag) |
| SC349 | *ubc13-Δ rad30Δ::hisG* VI(167260-167265)::(*lox66-3'lacZ-MET25*/lead) |
| SC350 | *ubc13-Δ rad30Δ::hisG* III(75494-75499)::(*lox66-3'lacZ-MET25*/lag) |
| SC351 | *ubc13-Δ rad30Δ::hisG* III(75494-75499)::(*lox66-3'lacZ-MET25*/lead) |

**Identification of TLS Products**

To identify the nucleotide inserted opposite the lesion site during TLS, a 1.5-kb fragment of the *lacZ* gene surrounding the lesion site was amplified by PCR using the oligos VP727 (5'- CATCCAGTGTCGAAAACGAG-3') and VP23 (5'- TTCTGCTTCAATCAGCGTGC-3'). The PCR fragments were then analyzed by HincII restriction endonuclease digestion, followed by gel analysis. PCR products that were not digested (error-prone TLS events) were sequenced to determine what kind of mutation had occurred. Table S2 shows the sequencing results.

**Table S2: TLS at CPD and TT(6-4) lesions**

Nucleotides inserted opposite the 5'-TTG-3' (TT is TT-CPD)

| | 3'-AAC-5' | 3'-AAG-5' | 3'-AAT-5' | 3'-AAA-5' | 3'-ATC-5' | 3'-AGC-5' | 3'-ACC-5' | 3'-AGG-5' | 3'-AGA-5' | 3'-CTC-5' | Other |
|---|---|---|---|---|---|---|---|---|---|---|---|
| WT | 82% | 7% | 0% | 11% | 0% | 0% | 0% | 0% | 0% | 0% | 0% |
| *rad30* | 80% | 7% | 0% | 7% | 3% | 0% | 0.5% | 0.5% | 0.5% | 0.5% | 0.5%[A] |
| *rev3 rev1* | 99.5% | 0% | 0% | 0% | 0% | 0.5%[B] | 0% | 0% | 0% | 0% | 0% |

[A] frameshift -G at +1
[B] and G instead of A at -1

Nucleotides inserted opposite the 5'-TTG-3' (TT is TT-(6-4))

| | 3'-AAC-5' | 3'-ATC-5' | 3'-AGC-5' | 3'-ACC-5' | 3'-AGA-5' | 3'-CAA-5' |
|---|---|---|---|---|---|---|
| WT | 70% | 4% | 23% | 1% | 1% | 1% |

**Comparison of different integration systems**

In order to test different integration strategies, we generated yeast recipient strains carrying 3′-end of *lacZ* fused to *lox66, loxJTZ17*, or *attB* integration site, and plasmids carrying LEU selective marker and the 5′-end of the *lacZ* gene fused to a *lox77, loxJT15*, or *attP* site, respectively. Following ectopic expression of Cre or phage phiC31 recombinase from a strong yeast promoter ADH (pKM4 and pKM5, containing TRP selection marker) we introduced the plasmids by electroporation. Cells were plated on selective indicator plates (SDa-LEU, X-gal). Plasmid pRS413 (containing HIS3 marker) was co-transformed and used as an internal control to normalize transformation efficiency between different vectors. Table S3 shows the efficiency of integration for each of the tested systems. The Cre/*lox66*/*lox71* recombination system shows the highest efficiency and is the one that we retained.

To evaluate if excision occurs with this system, several colonies were separately inoculated into SD-TRP medium and grown overnight in 30°C. Appropriate dilutions were plated on YPD plates, and the next day replica-plated on SDa-LEU plates. No colonies that lost the ability to grow without leucine were observed.

**Table S3:**

| Integrase | integration site | int. efficiency (CFU/ng DNA) |
|---|---|---|
| cre | *lox66/71* | 137 (±23) |
| cre | *loxJTZ17/JT15* | 34 (±5) |
| phiC31 | *attB/attP* | 9 (±4) |

**Protocol for integration of lesion containing vector by electroporation**

• Prior to integration, transform the chosen strain containing the chromosomal integration site with a plasmid carrying Cre recombinase (pKM34; contains TRP1 marker). Plate on SD agar -TRP plates with doxycycline (10 µg/ml) and incubate for 48h. Higher integration efficiencies are obtained with freshly transformed cells.

• Inoculate 10 ml of SD -TRP with doxycycline (5 µg/ml) until stationary phase ($OD_{600}$~3)

• Inoculate an aliquot of the overnight culture into 100 ml of prewarmed (30°C) YPD medium at a starting $OD_{600}$ about 0.3-0.4

• Continue to grow the cells at 30 °C on a platform shaker (225 rpm) until $OD_{600}$ is approximately 1.6

• Collect yeast cells by centrifugation at 3000 rpm for 3 minutes and remove the medium

• Wash the cell pellet twice with 50 ml ice-cold water and once with 50 ml of ice-cold electroporation buffer (1 M sorbitol / 1 mM $CaCl_2$)

• Condition the yeast cells by re-suspending the cell pellet in 20 ml conditioning buffer (0.1 M LiAc/10 mM DTT) and incubating for 30 min at 30 °C on a platform shaker (225 rpm)

• Collect the conditioned cells by centrifugation ad remove the medium. **Cells have to be kept on ice until electroporation**

• Wash once by 50 ml ice-cold electroporation buffer

• Resuspended the cell pellet in 100 - 200 µl electroporation buffer to reach a final volume of 1 ml. This is enough for 2 electroporations 400 µl each.

• Gently mix 400 µl of cells with DNA mix (100 ng of integrative plasmid with/without lesion, 100 ng of transformation control plasmid pRS413 containing HIS3 marker, and 12 µg of denatured carrier DNA) and transfer to a pre-chilled 0.2 cm electrode gap cuvette

• Keep on ice for 5 minutes until electroporation

• Electroporate at 2.5 kV/25 mF/ 800 Ω. Typical time constant ranges from 3.0 to 4.5 ms.

• Transfer electroporated cells into 6 ml of 1:1 mix of 1 M sorbitol : YPD Plus medium (ZymoResearch). Incubate at 30 °C on a platform shaker (225 rpm) for 1 hour

• Collect cells by centrifugation and resuspend in 5 ml 1 M sorbitol

• Plate appropriate dilutions on SD agar -LEU plates (with 80 mg/ml X-gal, 10 µg/ml doxycycline, and 1x phosphate buffer salts pH=7.00) and SD agar -HIS plates

• Incubate for at least 3 days at 30 °C, until colonies are large enough to count and blue color develops

**Electroporation buffer**

1 M sorbitol
1 mM $CaCl_2$

**10x phosphate buffer salts**
(add 100 ml to 900 ml SD agar)

39 g $NaH_2PO_4$•$2H_2O$
37 g $NaH_2PO_4$
$H_2O$ to 1 l
Adjust pH to 7.0 with NaOH

**Conditioning buffer**

0.1 M LiAc
0.2 10 mM DTT
